# Microglia target parvalbumin-expressing interneurons and tune gamma oscillations through P2Y12-dependent IL-1 receptor signaling

**DOI:** 10.64898/2026.11.25.769115

**Authors:** Ecem Tütüncü, Luna Šošo Zdravković, Michael Surala, Linus Alschäfscky, Niclas Gimber, Jan Schmoranzer, Julia Onken, Pawel Fidzinski, Jonas-Frederic Sauer, Jörg RP Geiger, Zoltan Gerevich, Christian Madry

## Abstract

Adaptation of neuronal network activity is critical for maintaining brain homeostasis and coping with changing computational demands. Microglia, the brain’s innate immune cells, are increasingly recognized to shape neuronal function in the healthy brain, yet their contribution to inhibitory circuit dynamics and network synchronization remains largely unexplored.

Here, we show that microglia establish a closed-loop feedback circuit that regulates synchronized network activity in the gamma frequency range, a key feature of cognitive processes that critically depends on fast-spiking parvalbumin-positive interneurons (PVIs). This mechanism operates in ex vivo mouse hippocampal and human neocortical circuits, indicating conservation across brain regions and species. During gamma oscillations, microglia preferentially recruit their processes to PVI somata through P2Y12 receptor-dependent purinergic signaling, whereas comparable targeting is absent around pyramidal neurons. Blockade of P2Y12 enhances gamma power, as does inhibition of interleukin-1 receptor type 1 (IL-1R1), an effect absent in P2Y12-deficient mice. P2Y12–IL-1R1 signaling functionally converges on adenosine A1 receptor activation, consistent with an IL-1β–adenosine-mediated neuroimmune feedback mechanism. At the cellular level, adenosine reduces excitatory synaptic input onto PVIs, whereas IL-1β attenuates their fast-spiking behavior.

Together, our findings establish microglia as physiological regulators of inhibitory network synchronization, linking preferential interactions with PVIs to P2Y12-coupled cytokine signaling and homeostatic gain control of gamma oscillations.

## Introduction

Microglia are increasingly recognized as key players that shape neural function in the developing and adult central nervous system to maintain brain homeostasis. Continuous movement of their highly motile processes is fundamental to the ability of microglia to survey the brain parenchyma (1, 2), thus safeguarding against damage and monitoring neuronal function. Although microglial dynamics appear random, their behavior is strongly influenced by neuronal activity (3–8), whereby microglia can sense both hypo- and hyperactivity of neurons, to which they respond by enhanced motility of their processes targeting neuronal structures (9, 10). Microglia can interact with neurons at various structural-functional levels, ranging from neurotransmission at individual synapses, direct cell-cell contacts to large-scale network regulation through the release of neuromodulatory substances (11).

Microglia-neuron interactions are particularly influenced by purinergic signaling driven by neuronal activity-dependent ATP release. Upon conversion to ADP by the microglia-enriched ectonucleotidase CD39 (12–14), ADP is sensed via high-affinity metabotropic P2Y12 purinoceptors which are exclusively and ubiquitously expressed by brain-resident microglia (15, 16). P2Y12 signaling plays a key role in establishing dynamic interactions with neurons, prominently occurring at specialized purinergic junctions formed between neuronal somatic regions and microglial processes (5, 6, 17). Microglia are preferentially attracted to highly active neurons, particularly during neuronal states close to the limits of physiological excitatory/inhibitory balance (4, 9, 17–19). Remarkably, enhanced microglial interaction can induce homeostatic feedback that attenuates neuronal firing and prevents transitioning into pathological activity states.

While most current knowledge relates to microglial interactions with principal neurons, comparatively little is known about how microglia regulate GABAergic interneurons and inhibitory network function under physiological conditions (19–21). Synchronized network activity in the 30–90 Hz gamma frequency range represents a hallmark of GABAergic neurotransmission and is fundamental to higher brain function including memory formation, cognition and sensory information processing (22). Gamma oscillations critically depend on the role of fast-spiking parvalbumin-positive interneurons (PVIs) (23, 24), which coordinate rhythmic activity of local pyramidal cell assemblies by providing perisomatic inhibition via fast synaptic feedback loops. Interestingly, microglia respond to specific rhythms of network activity within the gamma frequency range, inducing a complex neuroimmune signaling accompanied by transcriptional and functional adaptations (25–28). Moreover, pathology-associated microglial activation contributes to network dysfunction (29–31), suggesting that microglia are implicated in network synchronization. However, the cellular targets and signaling mechanisms through which microglia could modulate inhibitory circuit dynamics under physiological conditions remain elusive. Given their exceptionally high firing rates and central role in network synchronization (32, 33), PVIs represent a likely source of activity-dependent signals to microglia, thereby establishing homeostatic control at the network level.

To address these gaps in knowledge, we here investigated the role of microglia during gamma oscillations in mouse hippocampal and human neocortical circuits. Using electrophysiological, imaging, pharmacological, and genetic approaches, we examined how microglia respond to inhibitory network activity and interact with PVIs, integrating network, cellular and molecular levels. Our findings reveal a microglia-to-interneuron signaling pathway that mediates homeostatic control of gamma oscillations and establishes microglia as activity-dependent regulators of inhibitory network function.

## Results

### Gamma oscillations are sensed by microglial P2Y12 receptors

To investigate whether microglia respond to interneuron-driven network activity, we recorded local field potentials (LFPs) from the CA3 pyramidal cell layer in acute hippocampal brain slices from adult mice upon application of 150 nM kainate (KA) as an established model to trigger ex vivo gamma oscillations (34–36) (Fig. 1A-C). We first assessed microglial morphology as a sensitive readout of their responsiveness to changes in neuronal activity. In the presence of fully established gamma oscillations, which gradually emerge with increasing power (Fig. 1C), microglia displayed an increased number of processes with bulbous endings compared to non-oscillatory controls (Fig. 1D). These specialized terminal structures are associated with chemotactic and activity-dependent microglia-neuron interactions (4–6, 19), suggesting enhanced responsiveness of microglial processes during gamma activity. To determine whether these changes depend on ongoing neuronal activity, we acutely abrogated gamma oscillations by applying TTX. This interruption of network activity resulted in a marked decrease in both microglial ramification and bulbous endings compared to slices with sustained gamma oscillations (Fig. 1E).

**Figure 1.**
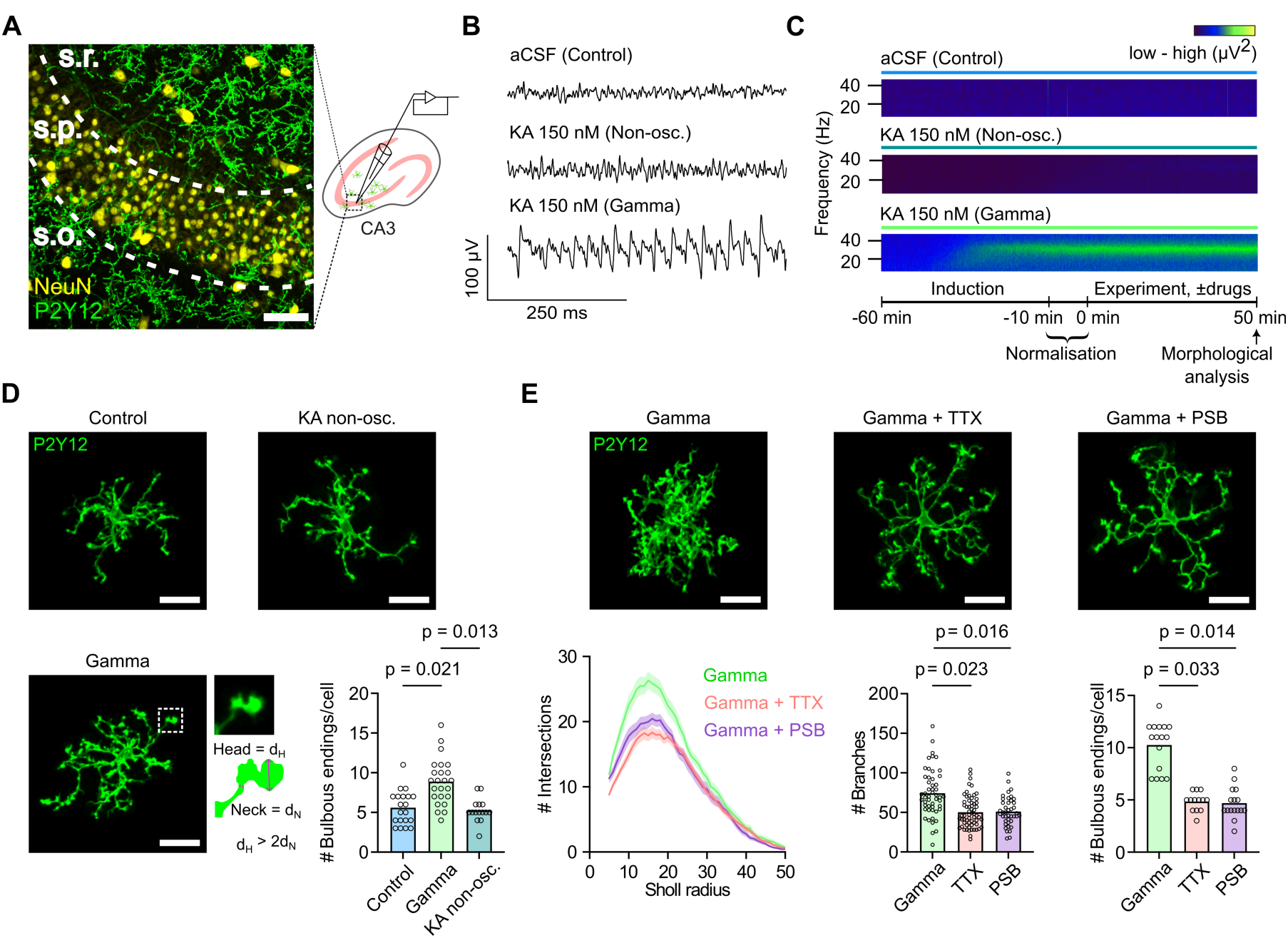
Gamma oscillations induce P2Y12 receptor-dependent reorganization of microglial processes. **A** Schematic of the experimental approach for local field potential (LFP) recordings in the hippocampal CA3 region. Left: Representative confocal image showing dense P2Y12^+^ microglial coverage throughout the CA3 pyramidal layer (NeuN^+^ neurons). Scale bar: 50 μm. **B**. LFP traces in aCSF and after application of KA, showing non-oscillating (Non-osc.) and gamma-oscillating (Gamma) responses. **C** Spectrograms showing the development of stable gamma oscillations following KA application compared with aCSF and KA non-osc. slices. Protocol of the application regime for electrophysiological and imaging experiments shown underneath. **D** Confocal images of individual microglia illustrating the increase in bulbous endings (defined as a head-to-neck diameter ratio >2) during gamma oscillations compared to non–oscillatory controls and analysis of changes (bottom right). Data points represent individual cells from 3 mice per condition; bars indicate the mean. P-values are from nested one-way ANOVA followed by Dunnett’s multiple comparisons tests. Scale bars: 20 µm. **E** Representative images showing reduced microglial process complexity (ramification) following blockade of gamma oscillations with 400 nM TTX or inhibition of P2Y12 receptors with 2 µM PSB-0739 (PSB) (top). Sholl analysis and quantification of the total number of branches and bulbous endings per cell (below). Data for Sholl analysis (intersections and branches) show mean ± SEM from 4 mice each for gamma and gamma + TTX, and 5 mice for gamma + PSB. Data for bulbous endings show mean from 5 mice each for gamma and gamma + PSB, and 4 mice for gamma + TTX. P-values are from nested one-way ANOVA followed by Dunnett’s multiple comparisons tests. Scale bars: 20 µm.

Since activity-dependent movement of microglial processes critically depends on P2Y12 receptors (5, 6, 16), we next tested whether similar effects could be observed after pharmacological inhibition of P2Y12 receptors using PSB-0739 (PSB) as a specific P2Y12 antagonist (37, 38). Blockade of P2Y12 receptors caused a reduction in microglial ramification and bulbous endings to a similar extent as TTX (Fig. 1E), suggesting that P2Y12 signaling mediates microglial process remodeling in response to activity-dependent purinergic signaling generated during gamma oscillations. Consistently, TTX failed to further alter microglial morphology in P2Y12-deficient mice, in which gamma oscillations were robustly elicited (Supplementary Fig. 1A-C). In addition, P2Y12 blockade without previous induction of gamma oscillations had no detectable effect on microglial ramification (Supplementary Fig. 1D), indicating engagement of P2Y12 receptors specifically during elevated neuronal activity. To distinguish gamma oscillation-associated microglial responses from effects of KA exposure alone, we further analyzed microglia in KA-treated slices in which gamma oscillations failed to emerge, under otherwise identical conditions. Notably, the increase in microglial bulbous endings was observed only in slices exhibiting gamma oscillations, but not in KA-treated slices lacking oscillatory activity (Fig. 1D). These findings indicate that microglial morphological changes are associated with the emergence of gamma oscillations rather than KA exposure alone. Together, our results demonstrate that microglia respond to gamma oscillations through a P2Y12-mediated mechanism, resulting in process remodeling and the formation of bulbous endings.

### Microglia selectively target PV^+^ interneurons during gamma oscillations via P2Y12 receptors

To examine the sites of microglial interactions during gamma oscillations, we focused on PV^+^ interneurons (PVI), which represent the principal fast-spiking interneuron population responsible for synchronizing neuronal activity during gamma oscillations, and are frequently contacted by microglia through specialized soma-targeting purinergic junctions (17). We therefore quantified the 3D radial distribution of P2Y12 immunofluorescence around the somata of CA3 PVIs, identified by PV-Cre-driven tdTomato expression, in slices after gamma oscillations or controls (Fig. 2A). Analysis of P2Y12 signal as a surrogate for perisomatic microglial coverage revealed a sharp increase beyond 5 µm from the center of the PVI soma, reaching a plateau indicating a constant distribution at distances greater than 20 µm. Consistent with the average dimensions of PVI somata, PV signal decayed within a radial distance of 5– 10 µm. Notably, PVI perisomatic microglial coverage was significantly increased in slices exhibiting gamma oscillations compared to non-oscillatory conditions, both in the absence and presence of KA (Fig. 2A, Supplementary Fig. 2A). Consistent with a P2Y12-dependent mechanism, inhibition of P2Y12 receptors abolished the gamma-associated enrichment of microglial processes around PVI somata, resulting in a radial profile comparable to non-gamma conditions (Fig. 2B). In contrast, differences in microglial abundance between gamma and non-gamma conditions were markedly reduced at greater distances from PVIs encompassing the subsequent 10 µm radial shell (Supplementary Fig. 2B), suggesting that gamma-associated microglial recruitment is concentrated in the immediate perisomatic region of PVIs. Interestingly, gamma oscillation-dependent recruitment of microglia was not observed around the somata of CA3 pyramidal neurons (Fig. 2C), despite an overall radial distribution of microglia similar to that observed for PVIs. Together, these findings demonstrate preferential P2Y12-dependent enrichment of microglial processes around PVI somata during gamma oscillations, an effect associated with oscillatory network activity rather than KA exposure alone.

**Figure 2.**
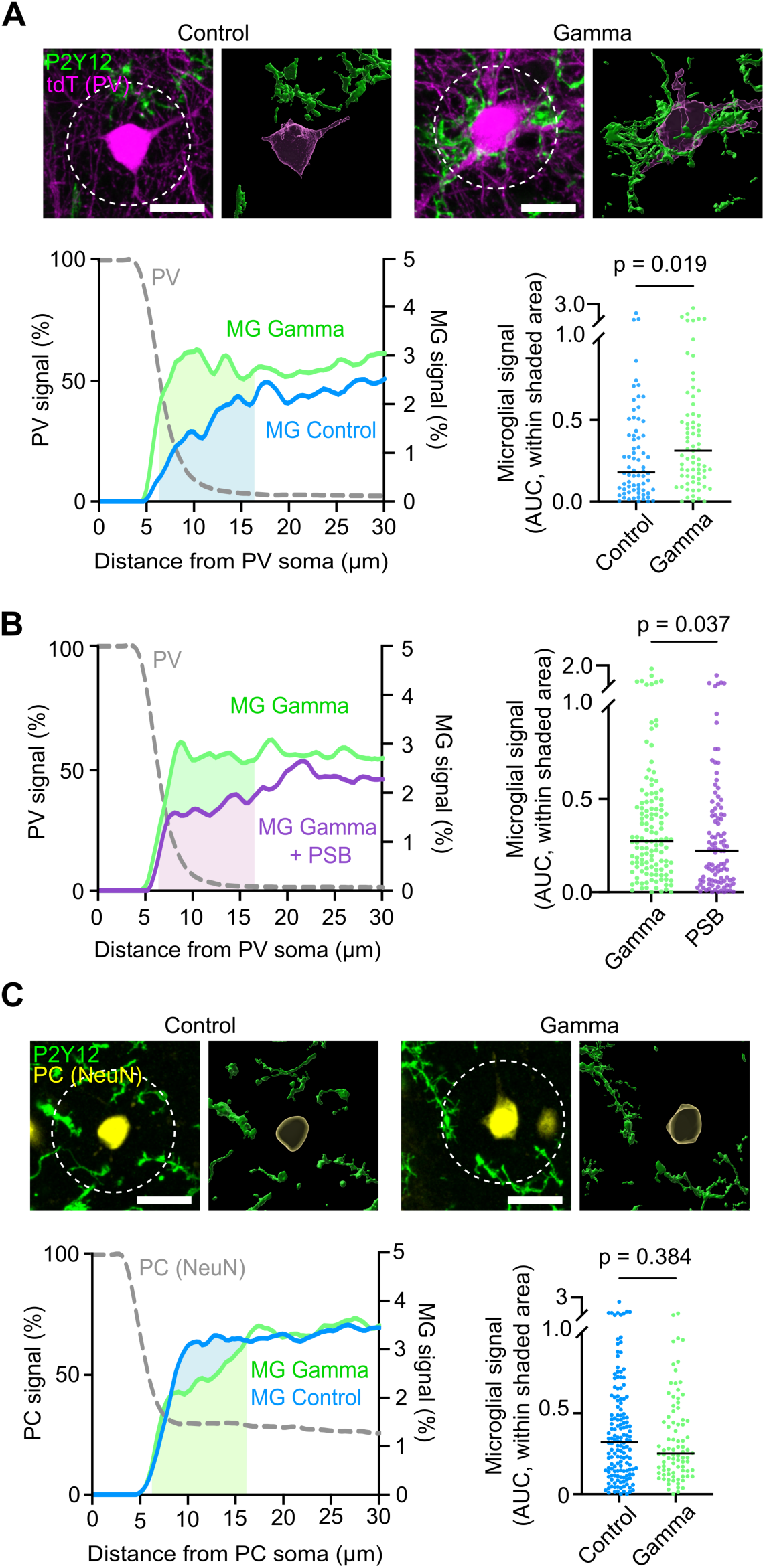
Microglia selectively target PV^+^ interneurons during gamma oscillations via P2Y12 receptors. **A** Top: Confocal and Imaris-rendered images showing the close apposition of microglial processes to PVI somata in the presence and absence of gamma oscillations. White circles indicate the extent of the spherical region used for radial analysis of the microglial signal. Scale bars: 20 μm. Bottom: Radial distribution of microglial P2Y12 fluorescence surrounding PVI somata (left). Microglial coverage quantified as the area under the fluorescence profile within the first 10 μm beyond the radius at which the PV signal had declined to 50% of its maximum (shaded area in the left graph) (right). **B** Left: Radial distribution of microglial P2Y12 fluorescence surrounding PVI somata during gamma oscillations with and without P2Y12 blockade (2 µM PSB). Right: Microglial coverage quantified as in **A**. Data points are individual soma measurements from n = 7 mice per condition; horizontal bars indicate the median. P-value is from Mann-Whitney test. **C** Same as in **A**, but for microglia surrounding pyramidal cells (PCs), defined as NeuN^+^PV^-^ neurons in stratum pyramidale. Scatterplots in (**A, C**) indicate individual soma measurements from 6 control and 5 gamma mice; horizontal bars indicate the median. P-values are from Mann-Whitney tests.

### P2Y12 signaling implements gain control of gamma oscillations in the mouse and human brain

We next examined whether P2Y12 signaling also regulates PVI-dependent oscillatory network activity. Acute blockade of P2Y12 receptors by PSB during established gamma oscillations led to a significant increase in gamma power over time, reaching an effect size of ∼ 30% relative to a 10-minute baseline before drug application (Fig. 3A-C). In contrast to gamma power, P2Y12 blockade produced only a minor change in oscillation frequency (Supplementary Fig. 3A). A comparable increase in gamma power was observed in P2Y12-deficient mice compared to littermate wild-type controls (Fig. 3D, E), whereas PSB failed to enhance gamma power in P2Y12-deficient mice (Supplementary Fig. 3B). Likewise, inhibition of P2X7 receptors using the selective antagonist A74003 (39, 40) did not alter gamma oscillations (Supplementary Fig. 3C), indicating that microglial regulation of gamma activity does not rely on purinergic signaling associated with high extracellular ATP concentrations required for P2X7 receptor activation.

**Figure 3.**
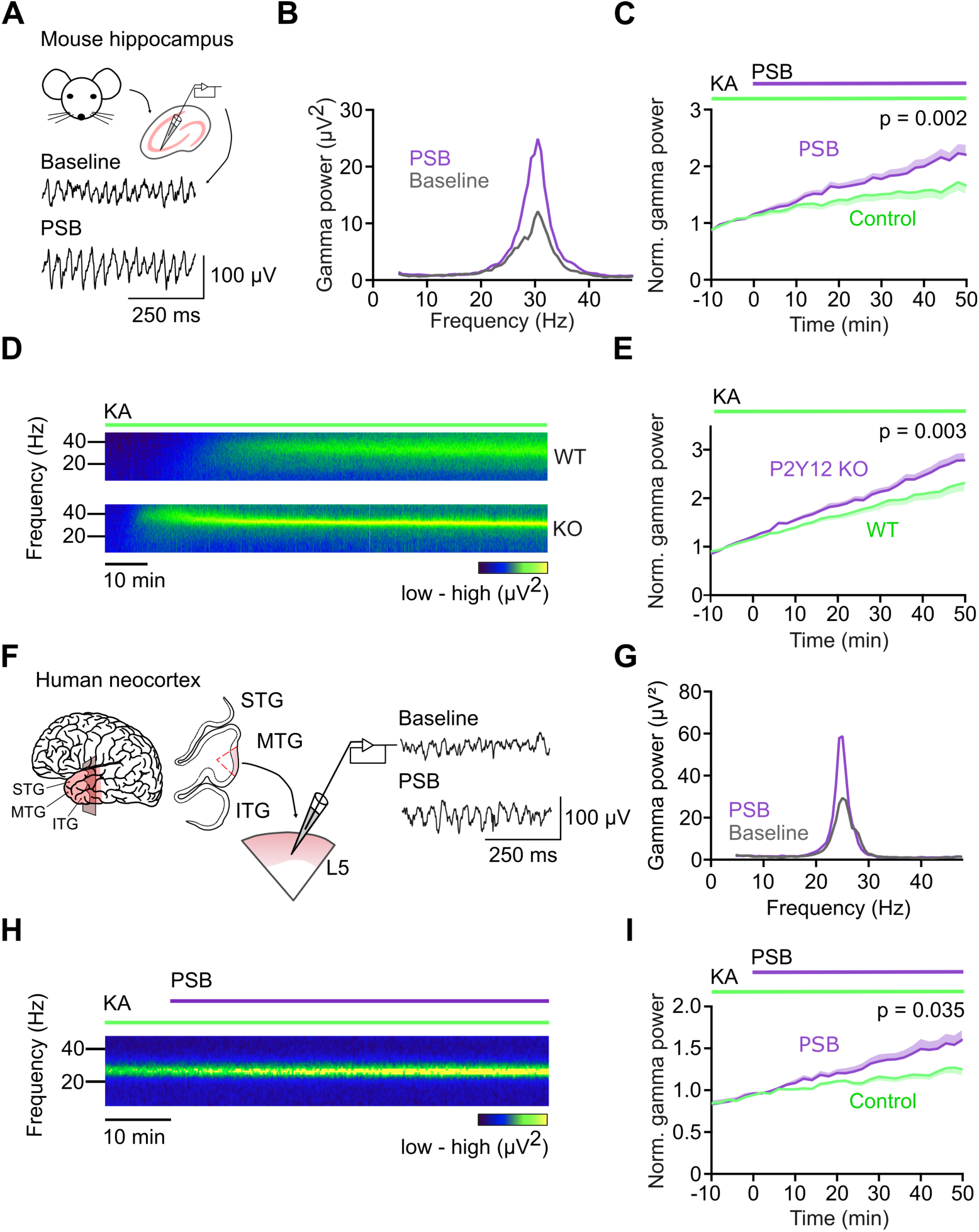
P2Y12 receptor blockade enhances gamma oscillation power in mouse hippocampus and human neocortex. **A** Representative LFP traces of gamma oscillations in the mouse CA3 region before (baseline) and after P2Y12 receptor blockade with 2 µM PSB-0739 (PSB). **B** Power spectra of recordings shown in **A**, illustrating increased gamma power following P2Y12 receptor blockade. **C** Normalized gamma power following PSB application after 60 min of KA perfusion and gamma stabilization. Gamma power was normalized to the 10 min interval preceding drug application. Data show mean ± SEM (n = 9 mice for control, n = 10 mice for PSB). Statistical significance was assessed using a linear mixed-effects model with an autoregressive AR(1) covariance structure including condition, time and their interaction. **D** Spectrograms of hippocampal gamma oscillations from wild-type (WT) and P2Y12-deficient (KO) mice, illustrating increased gamma power in KO mice. **E** Normalized gamma power in WT and P2Y12 KO mice. Normalization as in **C**. Data show mean ± SEM (n = 10 mice for WT, n = 15 mice for KO). Statistical analysis as in **C**. **F** Left: Schematic showing the preparation of acute cortical slices from human middle temporal gyrus (MTG) tissue. Right: Representative LFP traces during KA-induced gamma oscillations in layer 5 before and after P2Y12 receptor blockade with PSB. **G** Power spectra of human neocortical gamma oscillations before (baseline) and during application of PSB. **H** Spectrogram showing the change in established human neocortical gamma oscillations following application of PSB. **I** Normalized gamma power as in **C**, measured in human neocortex. Data show mean ± SEM (n = 4 patients for control, n = 5 patients for PSB). Statistical analysis as in **C**.

To assess translational relevance of this mechanism, we induced gamma oscillations in acute slices of human neocortical tissue resected during temporal lobe neurosurgery by application of 500 nM KA (Fig. 3F, G). Recordings were obtained from layer 5, a cortical region receiving dense perisomatic inhibitory input from PVIs (41) and exhibiting abundant microglial coverage (Supplementary Fig. 3D). Notably, inhibiting P2Y12 receptors also increased the power of human neocortical gamma oscillations (Fig. 3H, I), demonstrating a comparable onset and magnitude of effect as in mice. Together, these findings demonstrate that microglial P2Y12 signaling constrains gamma oscillation power in mouse hippocampal and human neocortical circuits, consistent with homeostatic regulation of oscillatory network gain.

### Microglial feedback on gamma oscillations depends on IL-1β signaling via engagement of IL-1 receptors

To investigate the molecular mechanisms underlying P2Y12-dependent regulation of gamma oscillations, we focused on microglial cytokine signaling, particularly IL-1β, which has been shown to be released from microglia in response to P2Y12 receptor activation (42, 43). Although IL-1β is generally regarded as a proinflammatory cytokine, at sub-inflammatory levels it acts as a potent neuromodulator in the healthy brain, regulating neuronal development, synaptic plasticity, and sleep-related processes (44, 45). We therefore tested whether gamma oscillations are affected upon disrupting signaling via IL-1 receptor type 1 (IL-1R1), the only signaling-competent receptor to which IL-1β binds (46, 47). Pharmacological blockade of IL-1R1 using Kineret (IL-1ra) as a highly selective antagonist caused a significant increase in gamma power, closely resembling the magnitude and time course observed following P2Y12 inhibition (Fig. 4A). Notably, this effect was absent in P2Y12 deficient mice (Fig. 4B). This pharmacological-genetic occlusion suggests a close functional coupling between P2Y12 and IL-1R1 signaling, consistent with IL-1R1 acting downstream of microglial purinergic activation.

**Figure 4.**
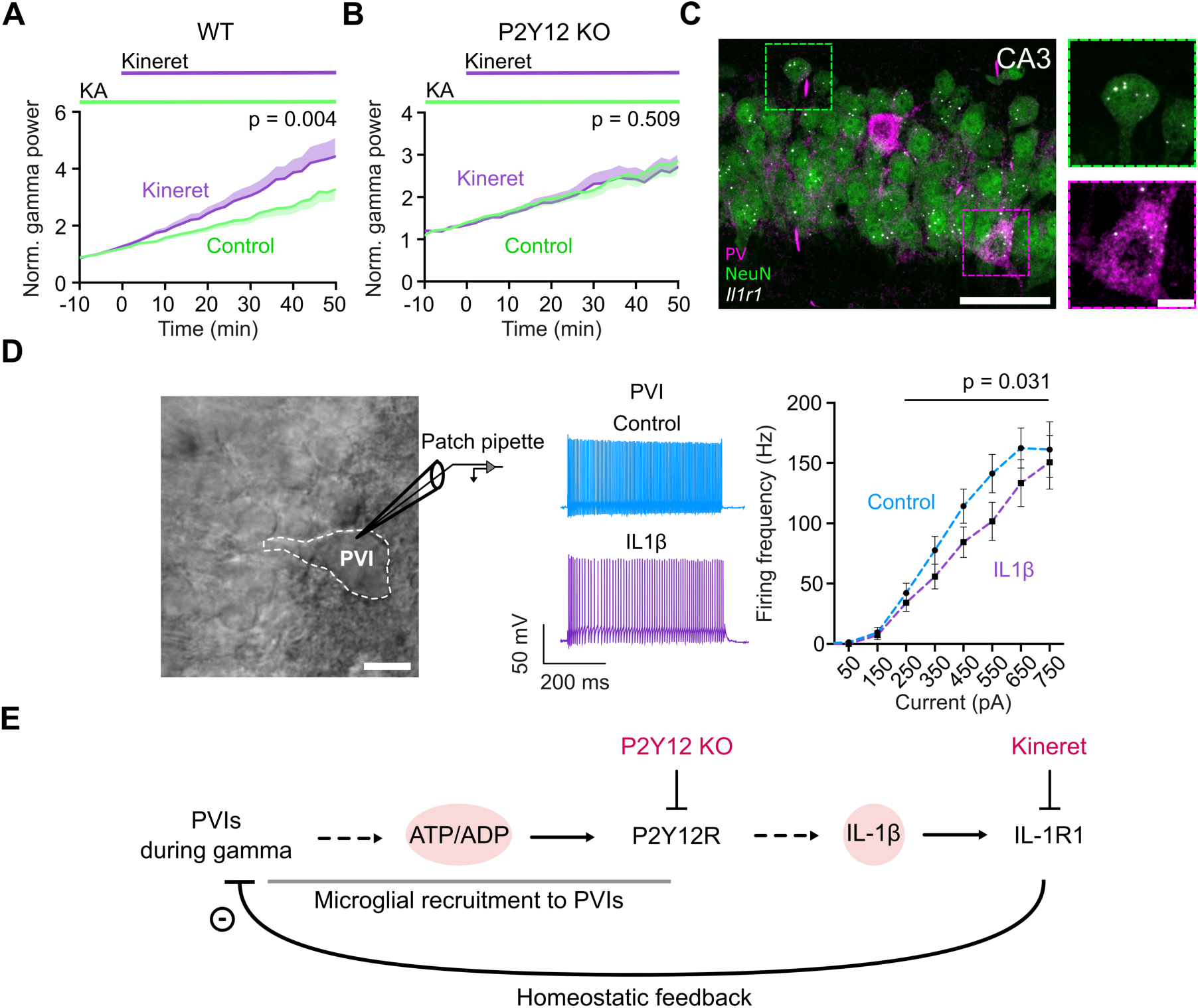
Microglial feedback on gamma oscillations depends on IL-1β–IL-1R1 signaling. **A** Time course of normalized gamma power before and after application of the IL-1 receptor type 1 (IL-1R1) antagonist Kineret (100 µg/ml). Kineret was applied after 60 min of KA perfusion, once stable gamma oscillations had been established. Gamma power was normalized to the 10 min interval preceding drug application. Data show mean ± SEM (n = 8 mice for control, n = 10 mice for Kineret). Statistical significance was assessed using a linear mixed-effects model (LMM) with an autoregressive AR(1) covariance structure including condition, time and their interaction. **B** IL-1R1 blockade by Kineret does not further enhance gamma power in P2Y12-deficient (KO) mice. Data show mean ± SEM (n = 4 mice per condition). Statistical significance was assessed using the same linear mixed-effects model as in **A**. **C** Left: Confocal image showing *Il1r1* mRNA detected by RNAscope in situ hybridization within NeuN^+^PV^-^ pyramidal cells and PV^+^ interneurons identified by immunofluorescence in the hippocampal CA3 region. Scale bar: 50 µm. Right: Close-up images showing *Il1r1* mRNA puncta within the soma of a pyramidal cell (top) and PVI (bottom). Scale bar: 10 µm. **D** Left: Differential interference contrast image of a patch-clamped PVI with its characteristic morphology and fast-spiking behavior (Control). Scale bar: 10 µm. Middle: Representative whole-cell current-clamp recordings from hippocampal PVIs showing action potential trains evoked by 550 pA current injection under control conditions (blue) and during application of 50 ng/mL IL-1β (purple). Right: Corresponding firing frequency-current relationship. Data show mean ± SEM (n = 12 PVIs from 5 mice). P-value is from LMM statistics with AR(1) covariance structure including condition, current and their interaction. **E** Mechanistic model illustrating gamma oscillation-evoked P2Y12-dependent purinergic signaling which induces a homeostatic feedback via IL-1R1 receptor activation.

To determine whether local IL-1β signaling associated with microglia-PVI interactions can directly affect PVI function, we examined the spatial expression of *Il1r1* mRNA in the hippocampus using RNAscope in situ hybridization. In addition to widespread *Il1r1* expression in NeuN^+^ CA3 pyramidal neurons and dentate gyrus granule cells, *Il1r1* transcripts are prominently expressed in PVIs in these regions, in contrast to other GABAergic interneuron types (Fig. 4C, Supplementary Fig. 4A, B). Consistently, whole-cell recordings from PVIs demonstrated that activation of IL-1R1 by IL-1β application reduced their excitability, as reflected by a lower firing frequency in response to increasing depolarizing current injection compared to control conditions (Fig. 4D). Together, these findings suggest that IL-1β contributes to P2Y12-dependent regulation of gamma oscillations and identify PVIs as potential cellular targets of IL-1β-mediated neuromodulation (Fig. 4E).

### IL-1R1-dependent regulation of gamma oscillations converges on adenosine A1 receptor signaling

To investigate how IL-1R1-dependent regulation affects gamma oscillations, we focused on adenosine signaling based on previous findings demonstrating that IL-1β can modulate hippocampal neurotransmission through an adenosine-dependent mechanism (44). Consistent with its role in hippocampal network activity (48, 49), application of adenosine markedly reduced gamma power, whereas blockade of A1 adenosine receptors by 8-CPT produced the opposite effect (Fig. 5A, C). To test whether adenosinergic signaling is implicated in the P2Y12–IL-1R1-dependent control of gamma power, we examined potential functional interactions between these pathways. Indeed, the increase in gamma power seen with P2Y12 blockade was completely abolished in the presence of either exogenously applied adenosine or the A1 receptor antagonist 8-CPT (Fig. 5B, D). Moreover, elevating extracellular adenosine fully prevented the increase in gamma power due to IL-1R1 inhibition (Fig. 5E). These findings reveal functional convergence of P2Y12, IL-1R1, and adenosine A1 receptor signaling in the regulation of gamma oscillations, consistent with IL-1R1 signaling acting downstream of microglial P2Y12 activation and upstream of adenosinergic feedback.

**Figure 5.**
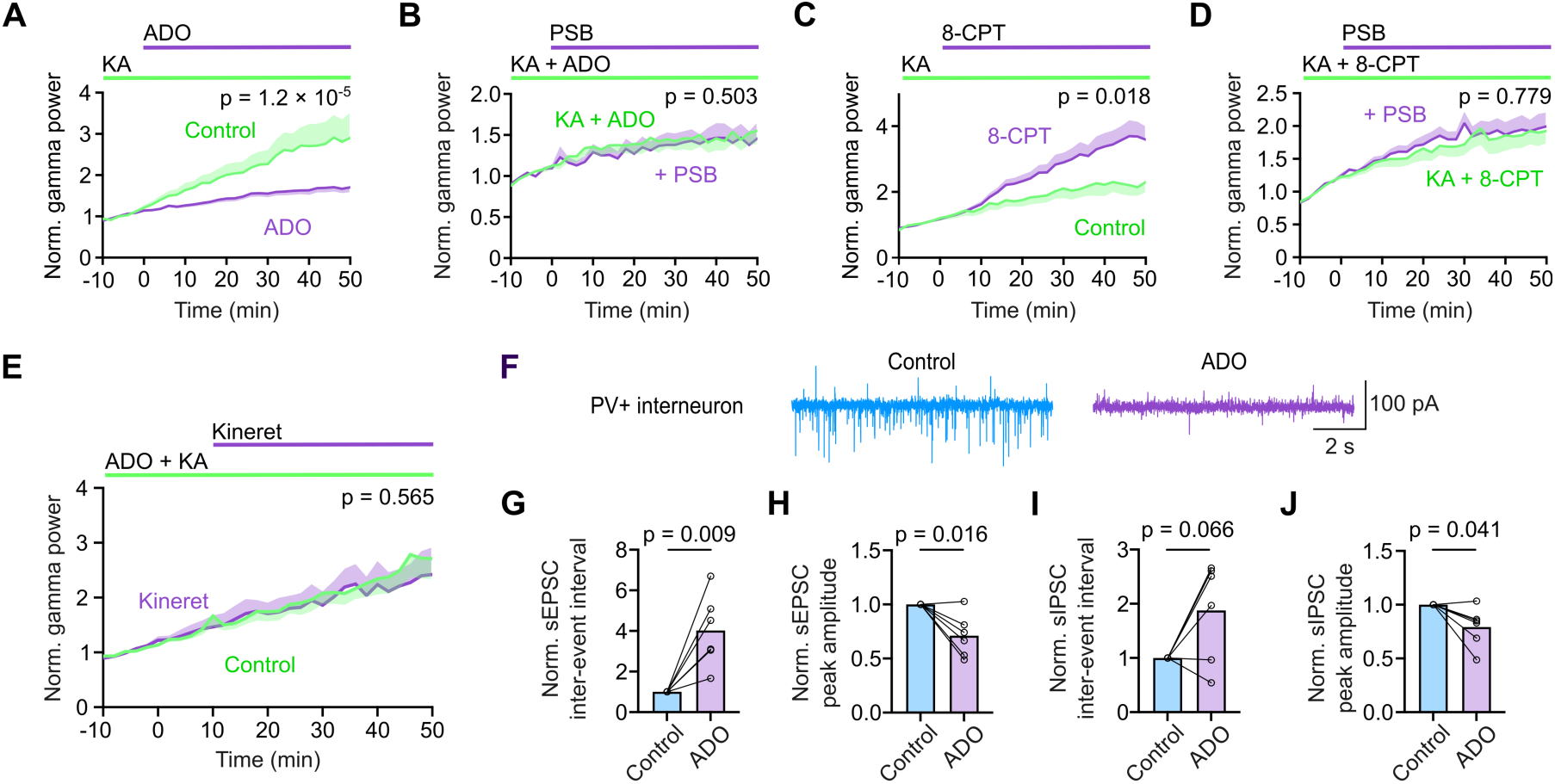
IL-1R1-dependent regulation of gamma oscillations converges on adenosine A1 receptor signaling. **A, C** Normalized gamma power before and after application of 50 µM adenosine (ADO) (**A**) or the A1 receptor antagonist 100 nM 8-CPT (**C**). Drugs were applied after 60 min of KA perfusion. Gamma power was normalized to the 10 min interval preceding drug application. Data show mean ± SEM (A: n = 5 mice for control and 10 mice for ADO; C: n = 4 mice for control and 12 mice for 8-CPT). Statistical significance was assessed using a linear mixed-effects model with an autoregressive AR(1) covariance structure including condition, time and their interaction. **B, D** P2Y12 receptor blockade with 2 µM PSB-0739 does not further enhance gamma power in the presence of 50 µM ADO (**B**) or 100 nM 8-CPT (**D**). Data show mean ± SEM (**B**: n = 6 mice per condition; **D**: n = 6 mice for control and 5 mice for PSB). Statistical analysis as in **A**. **E** IL-1R1 blockade by Kineret does not further enhance gamma power in the presence of 50 µM ADO. Data show mean ± SEM (n = 4 mice per condition). Statistical analysis as in **A**. **F** Representative recordings of spontaneous excitatory (sEPSC: downward deflections) and inhibitory (sIPSC: upward deflections) postsynaptic currents from voltage-clamped PVIs at -50 mV under control conditions (blue) and upon applying 50 µM ADO (purple). **G, H** ADO-dependent changes in sEPSC inter-event interval (**G**) and amplitude (**H**). Points represent individual PVI means over 5 min before (Control) and after 5 min of ADO application (n = 6 PVIs from 3 mice). Paired measurements are connected. P-values are from paired t-tests. **I, J** As in **G, H** but for ADO-induced changes in sIPSC inter-event interval (**I**) and amplitude (**J**) obtained from the same measurements.

We next investigated how adenosine affects PVI function at the cellular level. Previous studies have shown that adenosine suppresses the intrinsic excitability of hippocampal PVIs through A1 receptor activation (50). Since excitatory drive onto PVIs is a critical determinant of gamma oscillation generation and stability (34, 36), we asked whether adenosine also regulates their synaptic inputs. To address this, we performed whole-cell patch-clamp recordings from hippocampal PVIs to determine how elevated adenosine affects spontaneous excitatory and inhibitory postsynaptic currents (sEPSCs and sIPSCs; Fig. 5F). Application of adenosine significantly reduced both the frequency (i.e., increased inter-event intervals) and amplitude of sEPSCs (Fig. 5G, H), whereas changes in sIPSCs were more subtle (Fig. 5I, J). These findings suggest that adenosine acts primarily by reducing presynaptic glutamate release and excitatory input onto PVIs, which is expected to weaken the contribution of PVIs to neuronal synchronization during gamma oscillations (36). Together, these findings indicate functional convergence of P2Y12, IL-1R1, and adenosine A1 receptor signaling in the regulation of gamma oscillations, consistent with an IL-1R1-dependent adenosinergic feedback mechanism downstream of microglial P2Y12 activation.

## Discussion

In this study, we identify microglia as activity-dependent regulators of inhibitory network dynamics that constrain gamma oscillation power, bridging network, cellular and molecular levels. We show that microglia respond to oscillatory network activity through P2Y12 receptors and preferentially recruit their processes toward PVI somata. Functionally, P2Y12-dependent regulation of gamma oscillations is coupled to IL-1R1 signaling and converges on adenosine-mediated A1 receptor activation, consistent with a neuroimmune feedback mechanism involving IL-1β and adenosine. At the cellular level, adenosine reduces excitatory synaptic input onto PVIs, whereas IL-1β attenuates their fast-spiking behavior, identifying complementary signals that influence PVI-dependent inhibitory network synchronization (see summarizing scheme, Fig. 6).

**Figure 6.**
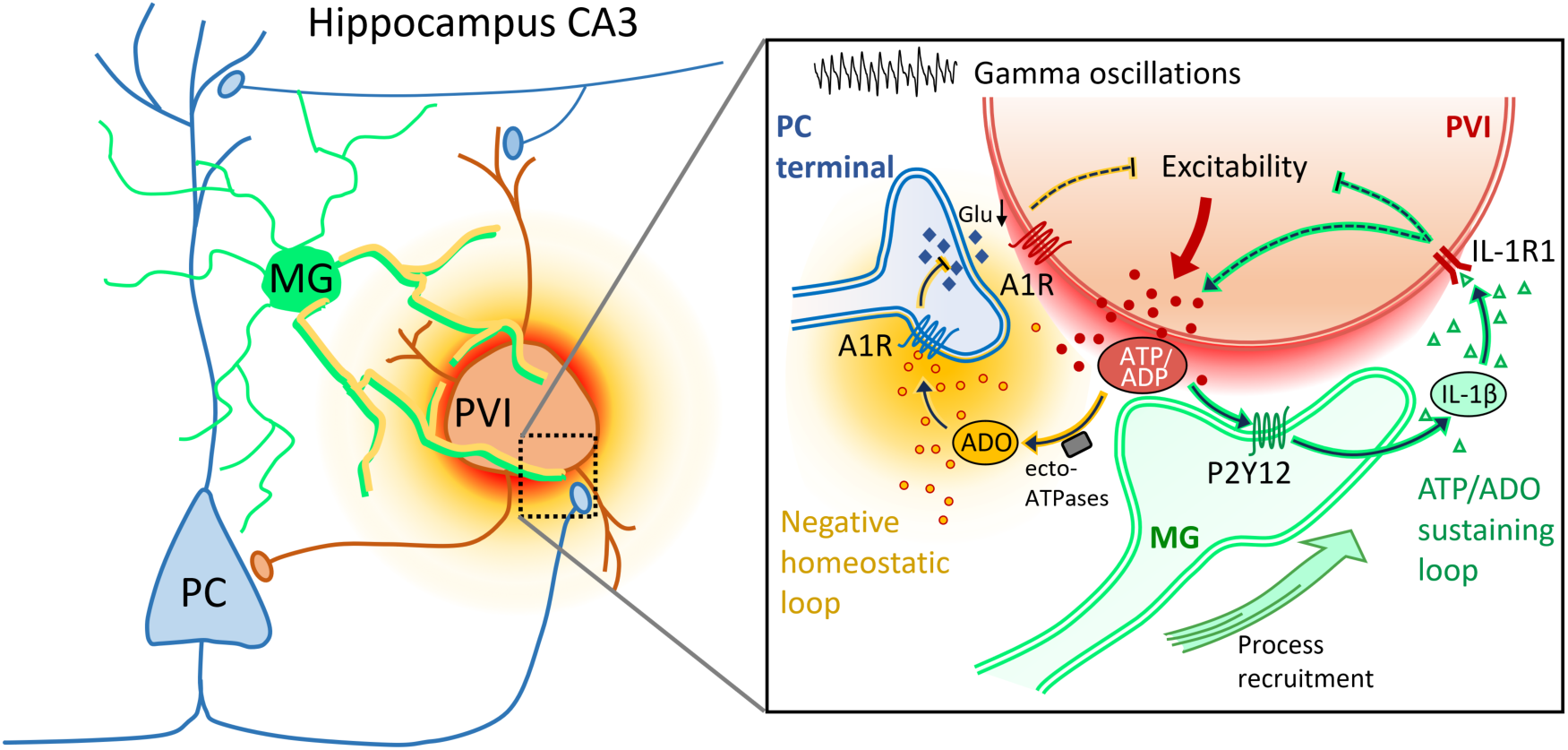
Proposed model of the mechanisms by which microglia regulate gamma oscillations. **Left:** Microcircuit view depicting the central role of PVIs in generating gamma oscillations by coordinating the rhythmic activity of local pyramidal cell assemblies via perisomatic feedback inhibition. During gamma, microglial processes (thick) are specifically recruited toward PVI somata in response to ATP release (red halo), generating an increase in adenosinergic tone (orange halo), thereby regulating gamma power. **Right:** Close-up view illustrating the proposed underlying mechanism. Gamma oscillation-dependent purinergic signaling (ATP/ADP) activates microglial P2Y12 receptors, promoting IL-1β release, which activates neuronal IL-1R1 and is proposed to facilitate extracellular ATP/adenosine generation, thereby enhancing local adenosinergic tone. The resulting increase in ADO tone (orange cloud) forms a negative homeostatic feedback loop (left) that limits gamma power via A1 receptor-mediated suppression of excitatory input onto PVIs. Although extracellular ADO may arise through enzymatic conversion of ATP, as illustrated, it may alternatively originate from direct neuronal ADO release (not shown).

Previous studies have established that microglia regulate neuronal activity and excitatory circuit function through purinergic signaling and adenosine-mediated negative feedback (4, 6, 10, 17, 18). Our findings extend this concept to the regulation of synchronized inhibitory network activity by linking preferential microglial interactions with fast-spiking PVIs to the control of gamma oscillation power. Importantly, we identify IL-1R1 signaling as a functionally coupled component of this mechanism, revealing an additional layer of neuroimmune control beyond previously described purinergic feedback pathways (18, 51, 52).

The ability of microglia to continuously survey the brain and interact with neurons is tightly regulated by purinergic signaling driven by neuronal activity-mediated ATP release (4–6). In this process, P2Y12 receptors act as high-affinity purinergic sensors that guide microglial processes toward highly active neurons and enable modulation of neuronal function. Purinergic signaling is also known to influence cortical and hippocampal gamma oscillations (49, 53), exerting both facilitatory and suppressive effects on network activity. Among these pathways, extracellular adenosine represents a particularly potent neuromodulator that suppresses gamma power via activation of high-affinity A1 receptors, whereas A2A receptor signaling can exert opposing effects (48, 49).

Similar to ATP release from excitatory neurons, P2Y12-dependent purinergic signaling acts as the central input pathway for microglial responses during gamma oscillation-associated neuronal activity, driving the targeted recruitment of microglial processes toward neuronal somata. Since ADP represents the preferred high-affinity ligand for P2Y12 receptors (38), ATP released from active neurons requires hydrolysis to ADP by ectonucleotidases such as CD39, which is highly enriched in microglia (13, 14). This local conversion is expected to effectively amplify P2Y12 signaling in the immediate vicinity of neuronal membranes, where the spatial range of ATP itself is highly restricted (54). During gamma oscillations, P2Y12-dependent recruitment of microglial processes occurs preferentially at PVI somata, suggesting enhanced local purinergic signaling at these sites. This is consistent with the formation of specialized purinergic junctions between microglial processes and neuronal somata, which are rapidly induced by neuronal activity, tightly linked to mitochondrial metabolism, and critically depend on P2Y12 receptor signaling (17), since such somatic junctions are formed not only with pyramidal neurons but also with PVIs. Given the exceptionally high firing rates and metabolic demands of PVIs, which necessitate elevated mitochondrial density and ATP turnover (55, 56), these interneurons are particularly suited to drive activity-dependent purinergic signaling. Given the rapid hydrolysis and limited diffusion of extracellular ATP compared to the greater stability and diffusion of adenosine (54, 57, 58), this conversion could translate a highly localized activity-dependent purinergic signal into a more spatially extended inhibitory feedback mechanism. Importantly, microglial process remodeling and preferential recruitment toward PVIs were associated with the emergence of gamma oscillations rather than KA exposure alone, supporting a context-specific response to oscillatory network activity.

A key finding of our study is the involvement of IL-1R1 signaling in microglial regulation of gamma oscillations. Pharmacological IL-1R1 blockade increased gamma power, closely resembling the effect of P2Y12 inhibition, whereas this response was absent in P2Y12-deficient mice. These findings reveal a functional coupling between microglial purinergic sensing and IL-1R1-dependent neuromodulation of synchronized network activity. Although IL-1β signaling has been extensively studied in pathological contexts associated with high extracellular ATP levels, accumulating evidence indicates that IL-1β also acts as a neuromodulator at much lower, physiological concentrations. Microglia can release IL-1β upon selective P2Y12 activation (42, 43), and IL-1β has been implicated physiologically in synaptic plasticity, learning, and sleep-wake regulation (46, 47, 59–61). The absence of an effect of P2X7 receptor inhibition in our experiments further supports the involvement of a purinergic signaling pathway distinct from classical P2X7-dependent inflammatory activation in microglial modulation of gamma power. Consistently, in vivo studies have demonstrated that neocortical microglia respond to visual stimulation-induced brain rhythms in the gamma-frequency range with specific morphological and functional changes (26, 28). This response is characterized by a frequency-specific cytokine profile that is distinct from classical inflammatory activation and reflects a homeostatic reaction to neuronal activity. Among the regulated factors, IL-1β emerges as one of the most strongly microglial-dependent cytokines induced by 40 Hz stimulation. These observations provide independent support for the notion that gamma-associated neuronal activity engages microglial cytokine signaling, in line with the involvement of IL-1R1 identified in our experiments.

Mechanistically, a close coupling between IL-1β signaling and adenosine-mediated modulation of synaptic transmission has been demonstrated in the hippocampus, where IL-1β triggers release of ATP and adenosine, thereby reducing CA3-CA1 neurotransmission (44, 45). Moreover, fractalkine signaling, a major neuron–microglia communication axis, engages Gi-coupled microglial CX3CR1 receptors to drive IL-1β-as well as adenosine-dependent modulation of synaptic plasticity (51, 60), paralleling Gi-mediated P2Y12 signaling. In healthy mice, IL-1 receptor expression is particularly prominent in the hippocampus (14, 62, 63), which is underscored by our finding of IL-1R1 expression not only in pyramidal neurons but also in PVIs. Consistently, at the cellular level, we find that adenosine and IL-1β exert complementary effects on PVI activity. Adenosine reduces excitatory synaptic input onto PVIs, whereas IL-1β attenuates their fast-spiking behavior, both of which are expected to reduce PVI recruitment and their contribution to rhythmic inhibitory network synchronization. Although adenosine also suppresses synaptic transmission and excitability in pyramidal neurons (64–66), our data indicate that modulation of PVIs represents a plausible cellular mechanism through which microglia influence gamma power.

Together with previous work demonstrating interactions between IL-1β and adenosine signaling in hippocampal circuits (44, 45), our findings support a model in which P2Y12-dependent microglial regulation of gamma oscillations involves IL-1R1 signaling and converges on adenosine A1 receptor activation. The pharmacological-genetic occlusion of gamma enhancement by IL-1R1 blockade in P2Y12-deficient mice, together with the absence of an additional effect of P2Y12 inhibition during A1 receptor blockade and the suppression of the Kineret response by exogenous adenosine, is consistent with IL-1R1 signaling functioning between microglial P2Y12 activation and adenosinergic feedback. Notably, these interventions predominantly affected gamma oscillation power, with comparatively minor changes in frequency, suggesting modulation of oscillatory network gain rather than disruption of the underlying rhythm-generating mechanisms (67).

Although our findings suggest a functional link between IL-1R1 signaling and adenosinergic feedback, the mechanisms underlying the regulation of extracellular adenosine tone remain incompletely understood (18, 45, 57, 66, 68). A purely ectonucleotidase-driven conversion of extracellular ATP to adenosine by microglia appears unlikely to fully account for this process, given the scarce expression of CD73 in cortical microglia (14, 69). The presence of IL-1R1 on both pyramidal cells and PVIs suggests that the regulation of extracellular adenosine involves neuronal IL-1R1 signaling. Adenosine may arise either from direct neuronal release (45, 65, 66, 70) or from extracellular ATP metabolism via ectonucleotidases and potentially additional cell types (18, 45, 54, 71). Thus, the cellular source of adenosine and the precise molecular mechanisms of its release remain to be established.

The identification of a microglia-driven feedback mechanism controlling gamma oscillations has potential implications for brain function and disease. Gamma oscillations are a key feature of cognitive processing, and their disruption is a hallmark of neurological and psychiatric disorders, including epilepsy, schizophrenia, and neurodegenerative diseases (22, 72–74). Given that P2Y12 is among the earliest homeostatic receptors downregulated during the transition to disease-associated microglial states (75, 76), our findings suggest that microglial activation may be accompanied by a progressive loss of mechanisms that normally sense and constrain neuronal network activity. Consistent with this idea, disruption of microglial Gi signaling induces network hyperexcitability (30), and loss of P2Y12 has been linked to increased epileptiform activity and seizure susceptibility, altered synaptic plasticity, and impaired neuronal survival following injury (6, 17, 77–79). Together, these observations suggest that disruption of microglial purinergic signaling may compromise mechanisms that normally stabilize neuronal network activity. Our demonstration that P2Y12-dependent modulation of gamma oscillations also operates in human neocortical tissue supports the potential relevance of this regulatory mechanism to human brain function. Understanding how microglial interactions with inhibitory circuits and their associated IL-1R1-dependent signaling are altered in disease may therefore provide new insights into network dysfunction in neurological and neurodegenerative disorders.

## Materials and Methods

### Animals and human brain tissue

Adult (12-16 weeks) C57BL/6J wild-type, constitutive P2Y12 KO (80), and PV-Cre-Ai9-tdTomato (Jackson laboratories) mice of both sexes were used. Mice were housed in individually ventilated cages under a 12 h light/dark cycle and ad libitum access to food and water. Procedures adhered to German animal welfare laws and were approved by the local health and social authorities (LaGeSo T0045/15, T-CH 0014/23, T-CH 0034/22).

Human neocortical tissue of the temporal lobe was obtained, with written informed consent, from five patients (4 males, 1 female; age range 27-59 years) undergoing surgery for drug-resistant epilepsy under approval of the Charité ethics committee (EA2/111/14). Brain tissue was transported from the operating theatre to the laboratory within 10-15 min in ice-cold sucrose-based solution as described previously (40) and was immediately subjected to slicing.

### Brain slice preparation

Mice: Acute horizontal hippocampal slices were prepared as described previously (40, 81). Briefly, after isoflurane anesthesia and decapitation, brains were removed and transferred to ice–cold carbogenated (95% O_2_ and 5% CO_2_) sucrose-based cutting solution (in mM): sucrose (75), NaCl (87), KCl (2.5), NaHCO₃ (25), NaH₂PO₄ (1.25), MgCl₂ (3), CaCl₂ (0.5), glucose (10). Final osmolarity was 310 to 315 mOsmol/L.

For LFP recordings, 400 µm slices were cut and transferred to custom-built interface chambers and perfused at 32-34 °C (1.6-1.8 ml/min) with carbogenated aCSF containing the following (in mM): NaCl (129), KCl (3), NaHCO₃ (21), NaH₂PO₄ (1.25), MgSO₄ (1.8), CaCl₂ (1.6) and glucose (10). Final osmolarity was 290 ± 5 mOsm/L. Slices for patch-clamp electrophysiology were prepared at 300 µm thickness and recovered for 15-20 min in cutting solution at 34 °C before storage at room temperature until experimental use.

Human material: After removal of the pia mater, human brain tissue was cut into 450 µm neocortical slices as previously described (40, 81) using ice-cold carbogenated sucrose-based solution as above. Following slicing, slices were allowed to recover for 30 min at 34°C in sucrose-based solution and were subsequently maintained in aCSF containing 4 mM KCl.

### Electrophysiology

Gamma oscillations were recorded from acute slices maintained under interface conditions (liquid-gas interface) with carbogenated aCSF at 32-34 °C and induced by the application of kainate (150 nM for mouse and 500 nM for human slices). Experiments were conducted on stable gamma oscillations 60 min after KA exposure. Local field potentials (LFPs) were recorded from the CA3 pyramidal layer with aCSF-filled electrodes (<10 MΩ), filtered at 0.1 Hz-1 kHz, and sampled at 5 kHz, using an EXT-02 B amplifier (npi) and a CED 1401 interface. Analysis of LFPs was done in Spike2 software. Power spectra were calculated in 120 s windows every 2 min; peak power, peak frequency, and half-bandwidth were extracted using a custom-written script. Gamma activity was defined by a peak frequency of 25–90 Hz and Q factor >0.5 (82).

Whole-cell patch-clamp recordings from visually identified PV^+^ basket cells (PVIs) were obtained in submerged slices perfused with carbogenated aCSF at 32-34 °C. Patch pipettes pulled from borosilicate glass had a tip resistance of 2.5–3.5 MΩ and were filled with an intracellular solution containing (in mM): K-gluconate (120), HEPES (10), EGTA (10), MgCl₂ (2), KCl (2.5), Na₂-phosphocreatine (5), Na₂-ATP (2) and Na₂-GTP (0.5). Series resistance (R_s_) was <20 MΩ and ∼ 60% compensated; recordings with a change in R_s_ of >20% were excluded. Signals were acquired using a MultiClamp 700B amplifier, filtered at 10 kHz, digitized at 20 kHz via a Digidata (both from Molecular Devices), and analyzed using pClamp10 software. Electrode junction potential was not compensated and was calculated to be 14.5 mV for the K-gluconate-based intracellular solution (determined using the LJP tool in pClamp10).

Excitability was assessed in current-clamp mode using 0.5 s current steps from -150 to 750 pA in 100 pA increments at the cells’ resting potential. Synaptic currents were recorded in voltage-clamped PVIs at -50 mV, allowing simultaneous analysis of sEPSCs and sIPSCs using template matching in Clampfit.

Application of drugs in electrophysiological experiments was via the bath perfusion. In time-matched control recordings without drug application, all measured parameters remained stable over the corresponding recording period. Pharmacological reagents and their sources are listed in Supplementary Table 1.

### Immunohistochemistry and RNAscope in situ hybridization

After LFP recordings, mouse slices were fixed overnight at 4°C in PFA in 0.1 M phosphate buffered saline (PBS; 0.9% NaCl), stored in PBS/0.05% NaN₃. To analyze microglial morphology in deeper tissue regions ∼ 100 µm below the slice surface, slices were re-sectioned from 400 µm to 80 µm thickness using a cryostat. Free-floating immunostaining was performed as described previously (81), with 72 hours primary antibody and overnight secondary antibody incubations at 4°C. Human neocortical slices were fixed in PFA for 4 h at 4°C immediately after slicing into 300 µm sections and subsequently processed using the same protocol, without re-sectioning. They were then treated with TrueBlackPlus according to the manufacturer’s instructions after immunostaining to reduce autofluorescence. Antibodies and dilutions are listed in Supplementary Table 1.

RNAscope Multiplex Fluorescent v2 with integrated co-detection was performed on 16-µm cryosections from PFA-fixed wild-type mouse brains according to the manufacturer’s protocol. *Il1r1* mRNA was detected using a mouse-specific probe with tyramide amplification and co-labeled for parvalbumin and NeuN to reveal PVIs. Slides were mounted with ProLong Gold and imaged using confocal microscopy. Reagents are listed in Supplementary Table 1.

### Imaging and morphometric analysis

Confocal Z-stacks spanning hippocampal CA3 strata oriens, pyramidale, and radiatum were acquired on a Nikon A1RSi+ microscope with a ×40 water-immersion objective (NA 1.25) at 0.35-µm intervals. Images had a resolution of 1024 by 1024 pixels and were processed using Fiji (ImageJ) by background subtraction, despeckling, 3D median filtering and intensity normalization.

For Sholl analysis, individual microglia from strata oriens, pyramidale and radiatum were reconstructed from Z-stacks using automatic Vaa3D tracing with manual correction where necessary (83). Number of branches and Sholl intersections were quantified using previously described MATLAB scripts (42).

For the analysis of bulbous endings, defined as protrusions with a head-to-neck diameter ratio >2 (19), individual microglia within strata oriens, pyramidale and radiatum were cropped and identified bulbous structures at their processes manually analyzed using Fiji (ImageJ).

For 3D radial analysis of microglia-PVI interactions, PV^+^ somata were segmented using a custom pipeline in Arivis Vision 4D software (https://github.com/ngimber/NeuroGlia3D) by morphological opening (1.5 µm), Gaussian smoothing (σ = 20 µm), Otsu thresholding, followed by a >400 µm³ volume cutoff. NeuN^+^ somata were segmented separately using Blob Finder (diameter 6 µm; threshold 8.36; watershed 29.5%; volume >100 µm³). NeuN^+^ profiles corresponding to PV^+^ somata or to cells located outside the stratum pyramidale were manually excluded. The remaining NeuN^+^PV^−^ somata were classified as putative pyramidal cells.

After extracting the center coordinates of all segmented somata, radial profiles were calculated using a custom Fiji (ImageJ) script (https://github.com/ngimber/RadialProfile3D). Microglial coverage within radial shells was quantified after binarization of the microglial P2Y12 immunofluorescence signal using Li’s minimum cross-entropy thresholding. The resulting 3D radial profiles were normalised to a range of 0-1 and averaged within two 10 µm radial distances bins (6.5-16.5 and 16.5-26.5 µm) from the soma center. The 6.5 µm inner boundary corresponded to the distance at which PV fluorescence declined to 50% of its maximum somatic signal.

All imaging and analyses were performed with the experimenter blind to condition and genotype.

### Statistics

Statistical analyses were performed using GraphPad Prism and IBM SPSS Statistics (version 31). To account for the hierarchical structure of the data and avoid pseudoreplication, nested statistical analyses and linear mixed-effects models were employed where appropriate, accounting for multiple observations obtained from individual brain slices and animals. Outliers were identified using ROUT (Q = 2%). Normally distributed data were analyzed using two-tailed nested t-tests or nested one-way ANOVA followed by Dunnett’s multiple comparisons tests. Non-normally distributed data were analyzed using Mann-Whitney U tests. Normalized gamma power and neuronal firing frequency were analyzed using linear mixed-effects models. Gamma power was sampled every 2 min and analyzed from 50–110 min, comprising 10 min before and 50 min after treatment onset. Time was centered at treatment onset (Time_Centered = Time − 60), with condition, Time_Centered, and their interaction as fixed effects, Mouse_ID as a random intercept, and an AR(1) covariance structure within Mouse_ID × Slice_ID. Firing frequency was analyzed across suprathreshold current injections ranging from 250 to 750 pA, with condition, injected current (modeled as a categorical factor), and their interaction included as fixed effects. Linear mixed-effects models were fitted using restricted maximum likelihood estimation with Satterthwaite approximation for degrees of freedom. Statistical significance was defined as p < 0.05.

## Abbreviations

aCSF: artificial cerebrospinal fluid
ADP: adenosine diphosphate
ADO: adenosine
ATP: adenosine triphosphate
IL-1β: interleukin-1 beta
IL-1R1: interleukin-1 receptor type 1
KA: kainate
LFP: local field potential
MTG: middle temporal gyrus
PSB: PSB-0739
PV: parvalbumin
PVI: parvalbumin-positive interneuron
P2Y12R: P2Y12 receptor
sEPSC: spontaneous excitatory postsynaptic current
sIPSC: spontaneous inhibitory postsynaptic current
TTX: tetrodotoxin
8-CPT: 8-cyclopentyltheophylline

## Declarations

### Ethics approval and consent to participate

All animal experiments were conducted in accordance with German animal welfare legislation and approved by the Berlin State Office for Health and Social Affairs (LaGeSo; T0045/15, T-CH 0014/23, T-CH 0034/22). The use of human temporal neocortical tissue was approved by the Ethics Committee of Charité – Universitätsmedizin Berlin (EA2/111/14). Written informed consent was obtained from all five patients undergoing surgery for drug-resistant epilepsy.

### Consent for publication

Not applicable.

### Availability of data and materials

The datasets used and/or analysed during the current study are available from the corresponding authors on reasonable request.

### Authors’ Contributions

Conceptualization: C.M., Z.G., E.T., J.R.P.G.; methodology: E.T., L.Š.Z., M.S., N.G., C.M., Z.G.; electrophysiological data acquisition and analysis: E.T., L.Š.Z., M.S., L.A.; imaging and quantitative analysis: E.T., L.Š.Z.; RNAscope experiments and image acquisition: M.S.; data visualization: E.T., L.Š.Z., M.S., Z.G.; resources: N.G., J.F.S., J.O., P.F.; funding acquisition: C.M., Z.G., J.R.P.G. (lead), E.T., L.Š.Z. (supporting); writing–original draft: C.M.; writing–review and editing: all authors; supervision: C.M., Z.G., J.R.P.G.

## Acknowledgments

We wish to thank Andrea Wilke for excellent technical assistance at the Institute of Neurophysiology of Charité Berlin. We are grateful to Ali Rifat, Michael Hadler and Henrik Alle for valuable comments and discussions on the manuscript. We also thank the Advanced Medical BioImaging Core Facility of the Charité – Universitätsmedizin Berlin (AMBIO) for support with the acquisition of the imaging data. We are grateful to the human brain research initiative of the Institute of Neurophysiology at Charité, in particular Henrik Alle, Franz Mittermaier, Henrike Planert and Michael Hadler, and associated institutions for general technical and logistical support. Moreover, we would like to thank Frank Konietschke at Charité for providing statistical support.

