## Supplementary Information for "Microglia target parvalbumin-expressing interneurons and tune gamma oscillations through P2Y12-dependent IL-1 receptor signaling"

**This PDF file includes:**

Supplementary Figures 1 to 4
Supplementary Table 1
Supplementary References

### Supplementary Figures

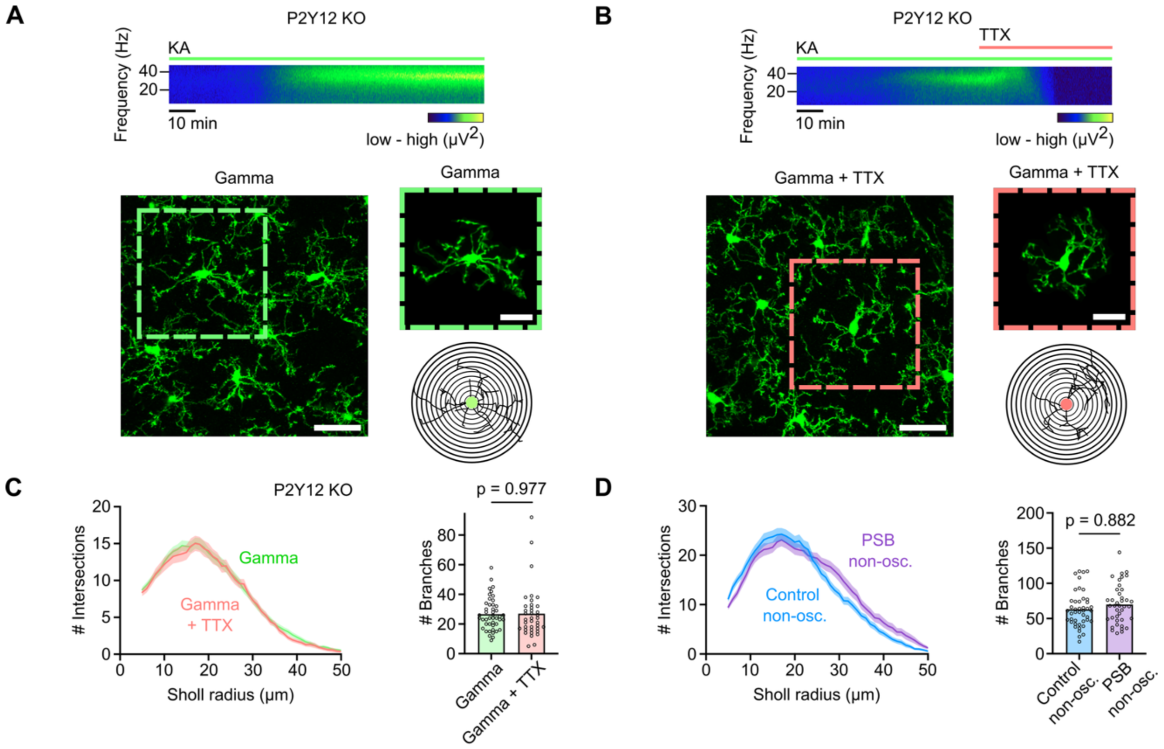


**Supplementary Fig. 1. Gamma oscillations drive microglial morphological changes.**

**A, B** Spectrograms and representative confocal images from CA3 stratum radiatum showing IBA1-stained microglia in hippocampal slices from P2Y12 KO mice in the presence of KA-induced gamma oscillations (Gamma) (**A**) and upon abrogating oscillations after application of 400 nM TTX (Gamma + TTX) (**B**). Scale bars: 30 μm. Smaller images show individual microglia and their morphological reconstruction for Sholl analysis. Scale bars: 20 μm.

**C** Sholl analysis showing the number of intersections (left) and total number of branches (right) for the conditions in **A** and **B** in P2Y12 KO mice. Data show mean ± SEM from n = 6 mice (gamma) and n = 4 mice (TTX). P-value is from two-tailed nested t-test.

**D** Sholl analysis showing the number of intersections (left) and total number of branches (right) of microglia in the absence of gamma oscillations (non-osc., no KA) without (control) and with application of PSB-0739. Data show mean ± SEM from n = 4 mice per condition. P-value is from two-tailed nested t-test.

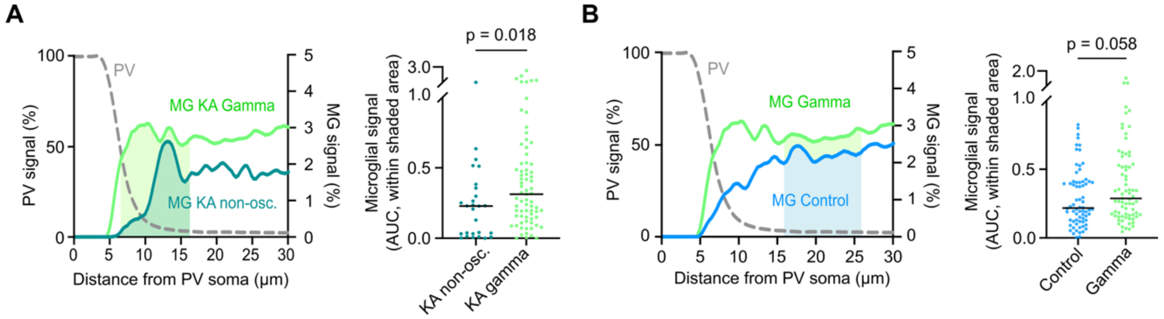


**Supplementary Fig. 2. Gamma oscillation-dependent microglia-PVI interactions and spatial distribution of microglia around PVI somata.**

**A** Left: Radial distribution of microglial P2Y12 fluorescence surrounding PVI somata for the indicated conditions. Right: Microglial coverage quantified as the area under the fluorescence profile within the first 10 µm beyond the radius at which the PV signal had declined to 50% of its maximum. KA-induced gamma oscillations, but not exposure to KA without developing gamma oscillations (KA non-osc.), increased microglial coverage around PVI somata. Data points are individual soma measurements from n = 3 mice for KA non-osc. and n = 7 mice for gamma; horizontal bars indicate the median. P-value is from Mann-Whitney test.

**B** Left: Radial distribution of microglial P2Y12 fluorescence surrounding PVI somata. Right: Microglial coverage at greater distances from the PVI soma, quantified as the area under the fluorescence profile within the subsequent 10 µm radial shell beyond the proximal region analyzed in Fig. 2A. Data points are individual soma measurements from n = 6 mice for control and n = 5 mice for gamma; horizontal bars indicate the median. P-value is from Mann-Whitney test.

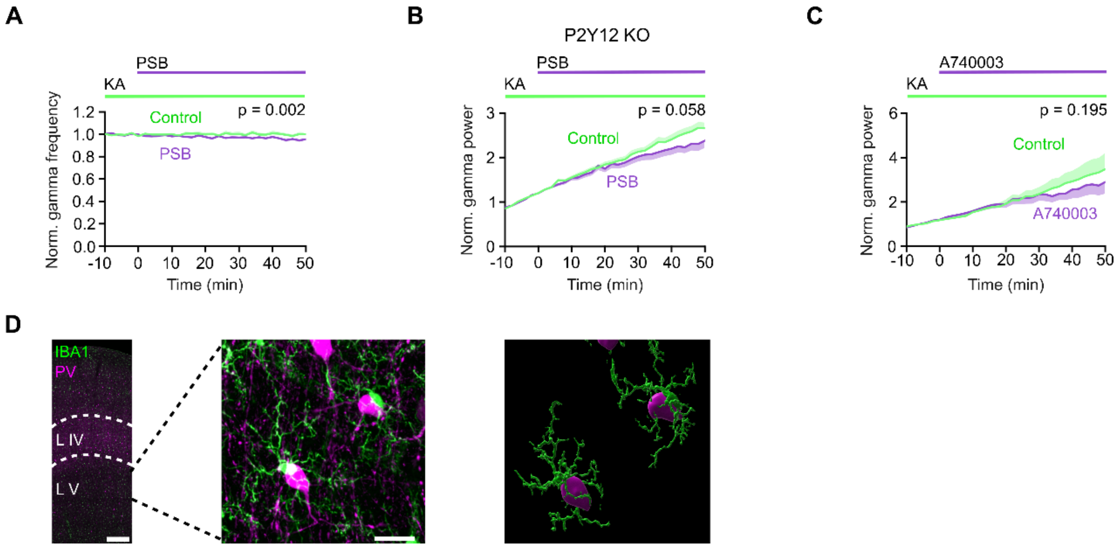


**Supplementary Fig. 3. Effects of microglial purinergic signaling on gamma oscillations.**

**A** Time course of normalized gamma frequency before and after P2Y12 receptor blockade with 2 µM PSB-0739 (PSB). Data show mean ± SEM (n = 9 mice for control, n = 10 mice for PSB). Statistical significance was assessed using a linear mixed-effects model with an autoregressive AR(1) covariance structure including condition, time, and their interaction (see Methods).

**B** Time course of normalized gamma power before and after P2Y12 receptor blockade in P2Y12-deficient (KO) mice. Data show mean ± SEM (n = 15 mice for control, n = 13 mice for PSB). Statistical significance was assessed using the same linear mixed-effects model as in **A**.

**C** Time course of normalized gamma power before and after P2X7 receptor blockade with 20 µM A740003. Data show mean ± SEM (n = 6 mice per condition). Statistical significance was assessed using the same linear mixed-effects model as in **A**.

**D** Confocal images showing microglia (IBA1, green) and parvalbumin-expressing interneurons (PV, magenta) in human neocortex. Left: Overview illustrating cortical layers. Middle: Layer V micrograph. Right: Segmentation showing microglial morphology and proximity to PV-positive interneurons. Scale bars: 500 µm (left), 20 µm (middle).

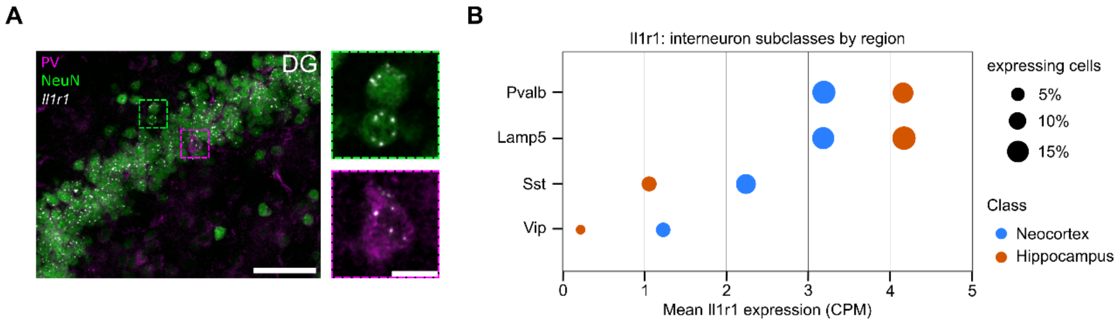


**Supplementary Fig. 4. IL-1 receptor expression in hippocampal and neocortical PVIs.**

**A** Left: Confocal image showing *Il1r1* mRNA detected by RNAscope in situ hybridization within $NeuN^{+}PV^{-}$ granule cells and $PV^{+}$ interneurons identified by immunofluorescence in the hippocampal dentate gyrus. Scale bar: 50 µm. Right: Close-up images showing *Il1r1* mRNA puncta within the soma of a granule cell (top) and PVI (bottom). Scale bar: 10 µm.

**B** Expression profile of *Il1r1* mRNA across GABAergic interneuron populations in the hippocampus and neocortex of adult mice. Single-cell RNA-sequencing data were obtained from the Allen Brain Cell Atlas (Mouse Whole Cortex and Hippocampus SMART-seq v4), Allen Institute for Brain Science (1). Pvalb, parvalbumin; Lamp5, lysosomal-associated membrane protein family member 5, representing predominantly neurogliaform cells; Sst, somatostatin; Vip, vasoactive intestinal peptide. CPM, counts per million.

**Tables**

**Supplementary Table 1. Reagents and resources**

| Reagent / Resource | Supplier | Catalogue no. | Dilution / Use |
|---|---|---|---|
| **Chemicals and recombinant proteins** | | | |
| Kainic acid (KA) | Tocris Bioscience (Bio-Techne) | 222 | 150 nM (mouse)<br>500 nM (human) |
| PSB-0739 | Tocris Bioscience (Bio-Techne) | 3983 | As indicated |
| A740003 | Tocris Bioscience (Bio-Techne) | 3701 | As indicated |
| 8-Cyclopentyl-1,3-dimethylxanthine (8-CPT) | Tocris Bioscience (Bio-Techne) | 6137 | As indicated |
| Adenosine | Sigma-Aldrich | A9251 | As indicated |
| Mouse IL-1β | Biomatik | RPU42172 | As indicated |
| Tetrodotoxin citrate (TTX) | Hello Bio | HB1035 | As indicated |
| Kineret® (anakinra) | Sobi Pharmaceuticals | — | As indicated |
| **Antibodies and reagents for immunohistochemistry** | | | |
| Rabbit anti-P2Y12 | AnaSpec | AS-55043A | 1:1000 |
| Rabbit anti-Iba1 | Synaptic Systems | 234 003 | 1:1000 |
| Guinea pig anti-Iba1 | Synaptic Systems | 234 308 | 1:500 |
| Guinea pig anti-NeuN | Merck/Millipore | ABN90 | 1:500 |
| Rabbit anti-NeuN | Merck/Millipore | ABN78 | 1:200 |
| Guinea pig anti-PV | Swant | GP72 | 1:200 |
| Mouse anti-PV | Swant | PV235 | 1:5000 |
| Goat anti-rabbit Alexa Fluor 488 | Thermo Fisher Scientific | A-11008 | 1:1000 |
| Goat anti-guinea pig Alexa Fluor 633 | Thermo Fisher Scientific | A-21105 | 1:500 |
| Goat anti-guinea pig Alexa Fluor 488 | Invitrogen | A-11073 | 1:500 |
| Goat anti-mouse Alexa Fluor 555 | Invitrogen | A-21422 | 1:500 |
| DAPI-containing mounting medium | Abcam | ab104139 | — |

| | | | |
|---|---|---|---|
| TrueBlack Plus Lipofuscin Autofluorescence Quencher | Biotium | 23014 | 1× |
| ProLong Gold | Invitrogen | P36934 | — |
| **In situ hybridization** | | | |
| RNAscope Multiplex Fluorescent Reagent Kit v2 | Advanced Cell Diagnostics (Bio-Techne) | — | Manufacturer's protocol |
| Mouse *Il1r1* probe | Advanced Cell Diagnostics (Bio-Techne) | — | Manufacturer's protocol |
| **Experimental models** | | | |
| PV-Cre mice | The Jackson Laboratory | 008069 | — |
| Ai9 reporter mice | The Jackson Laboratory | 007909 | — |
| **Software** | | | |
| MATLAB | MathWorks | https://www.mathworks.com/ | |
| pClamp10 | Molecular Devices | https://www.moleculardevices.com/ | |
| Spike2 | Cambridge Electronics | https://ced.co.uk/ | |
| Vaa3D | HHMI Janelia Research Campus Allen Institute for Brain Science | http://www.vaa3d.org | |
| Imaris 10.0.0/10.1.1 | Oxford Instruments | https://imaris.oxinst.com/ | |
| FIJI/ImageJ | National Institutes of Health (NIH) | https://imagej.nih.gov/ | |
| Arivis Vision 4D | Carl Zeiss Microscopy Software | https://www.arivis.com/ | |
| GraphPad Prism | GraphPad | https://www.graphpad.com/ | |
| SPSS Statistics v31 | IBM | https://www.ibm.com/ | |

**Supplementary Material References**


1. Z. Yao, *et al.*, A high-resolution transcriptomic and spatial atlas of cell types in the whole mouse brain. *Nature* **624**, 317–332 (2023).
